# Analysis of Naturally Arising Simple Deletions in Single Human Colon Crypts with Age

**DOI:** 10.64898/2026.09.30.755170

**Authors:** Yong Hwee Eddie Loh, Michael R. Lieber, Timothy Okitsu, Cindy Okitsu, Jordan Wlodarczyk, Zarko Manojlovic, Chih-Lin Hsieh

**Affiliations:** Norris Medical Library, University of Southern California, Los Angeles, CA, USA; Department of Pathology, University of Southern California, Los Angeles, CA, USA; Department of Urology, University of Southern California, Los Angeles, CA, USA; Department of Surgery, University of Southern California, Los Angeles, CA, USA; current affiliation: Intermountain Health Saint Joseph Hospital, Denver, CO, USA

**Author notes:** Contributed equally (Yong Hwee Eddie Loh, Michael R. Lieber).

## Abstract

Changes of DNA sequence in the human genome may contribute to human aging, but it has been difficult for the field to define the details of the contribution. Cancer cells have altered ratios of DNA damage and repair pathways, and thus it is inappropriate to utilize them as models for normal cells. Also, somatic mutations from each lineage cannot be clearly identified in cell populations with multiple lineages.

Normal human colon crypts are one of the few mini-organ systems where monoclonal cell populations can be found for somatic mutation analysis. In a parallel study, we described complex insertion/deletion (indel) events in normal human colon. Here we focus on simple deletions that lack any nucleotide addition or adjacent base substitutions. We find that ∼58% of the simple deletions are consistent with a strand slippage event, and the remainder provide insights into other important repair processes. Among the 4,264 ≥2-nucleotide simple deletions, ∼71% are most consistent with nonhomologous DNA end joining (NHEJ), with <1.5% of the events being more consistent with alternative end joining (aEJ). The incidence of NHEJ-type simple deletions is ∼8-fold more frequent than the complex indels that also show NHEJ features. Similar to the complex indels, the simple deletions show a linear trend of event accumulation (positive slope) with an overall ∼10-fold increase in absolute number across the human lifespan, 0.8 months to 90 years old, for individuals with no radiation or chemotherapy treatment. Interestingly, radiation treatment increases non-slippage deletions with little effect on slip deletions.

## Introduction

Beyond simple point mutations, natural occurring changes in genomic DNA can arise via several mechanisms. Replicative DNA synthesis can result in strand slippage events, which can give rise to insertions or deletions, especially in repetitive sequence regions ^[1]^. Naturally arising DNA damage from background ionizing radiation and oxidative metabolism can lead to double-strand breaks (DSBs) in the non-repetitive regions ^[2]^. There are four repair pathways for DSB in eukaryotic cells ^[3]^. The two main pathways are homologous recombination (HR) and nonhomologous DNA end joining (NHEJ). HR occurs predominantly during S-phase, and NHEJ takes place predominantly during the remainder of the cell cycle ^[4]^. Single-strand annealing (SSA) and alternative end joining (a-EJ) are two additional pathways that function primarily during DNA synthesis. HR and SSA use various lengths of strong sequence homology and can be readily distinguished from NHEJ ^[5, 6]^. The a-EJ pathway is primarily involved in repairing collapsed replication forks and relies heavily on DNA polymerase theta (Pol Q) ^[6, 7]^. The Pol Q transcript and protein levels are low in human colon epithelium, limiting the role of the a-EJ pathway in repairing DNA damage in this tissue^[8]^.

In normal individuals, the components of the NHEJ machinery are constitutively expressed in all cells of the body at substantial levels ^[5, 9]^. The major components include Ku, DNA-PKcs, Artemis, polymerases mu and lambda, DNA ligase IV, XRCC4, and XLF ^[3]^. The mechanistic steps and roles of the components are discussed elsewhere ^[4]^. The structural biology underpinning the mechanistic steps has been elucidated ^[10]^. NHEJ is distinctive in that it does not require any DNA sequence homology or even microhomology (MH), and that entirely incompatible DNA ends can be joined. When microhomology is present, as is often pre-determined in experimental systems, then the MH stabilizes the alignment of the two DNA ends in NHEJ, thereby favoring the outcome toward MH usage ^[4, 5]^. Polymerases mu and lambda are able to add nucleotides in either a template-independent or in a template-dependent manner ^[5, 11]^. Cellular studies of NHEJ have utilized various systems and approaches. Integrated target genes can be selected for event outcomes that permit drug selection of the target gene ^[12, 13]^. Other studies have transfected linearized plasmids or DNA fragments followed by recovery ^[14, 15]^. Importantly, nearly all these studies rely on cell lines in culture in which integration of target genes can be readily achieved. One excellent mammalian cell study used organoids grown from single cells isolated from disaggregated human colon crypts. However, this required DNA extraction after ≥2 weeks of growth in a 20% oxygen tissue culture incubator, and mutations arising during the initial days of growth at non-physiologic oxygen cannot be ruled out ^[16]^.

Conventional whole genome sequencing (WGS) can obtain the consensus sequence of all the cells in a sample cell population. However, heterogeneity of the DNA sequence in a non-clonal cell population makes reliable detection of DNA damaged site repair extremely difficult as discussed previously ^[1]^. For the first time, naturally arising somatic mutations can be comprehensively examined directly without DNA extraction due to our technical breakthrough in generating high-depth and high-quality DNA sequences from natural cell clones, such as single human colon crypts, ^[17, 18]^.

Based on the findings from biochemical and cellular studies, NHEJ events often appear to involve a small deletion with misincorporation of nucleotides. We termed these as complex indels and developed a customized algorithm to identify and analyze these ^[19, 20]^. In our study here, we examine simple deletions that are not complicated by adjacent sequence changes, such as insertions and base changes. The largest category of deletion events occurs in repetitive regions and accounts for 58.4% (49.1% are 1-nucleotide deletions and 9% are ≥2-nucleotide deletions) of the total set of simple deletions. The second largest category (23.7% of the total) consists of deletions ≥2-nucleotide that have no hallmarks of slippage, and these appear to be repaired by NHEJ. Interestingly, the number of NHEJ-type simple deletions have a very similar age-dependence as the age-dependence of the complex indels. We discuss the mechanistic aspects and functional impact of these findings.

## MATERIALS AND METHODS

### Tissue collection, crypt isolation, sequencing library construction, and whole-genome sequencing

As previously described, normal colon tissue was collected through the Pathology Department Tissue Procurement Core with an approved IRB from individuals who have undergone surgery to remove part of the colon under the standard of care at either Keck Hospital of USC or Children’s Hospital Los Angeles. Tissue processing, crypt isolation, and WGS library construction have been described previously ^[17]^.

A total of 106 crypt libraries and 21 bulk libraries are constructed from 21 individuals of age 10 months to 90 years old. After quality control, libraries are pooled and sequenced on S4 flow cells using NovaSeq 6000 (Illumina, San Diego) S4 300 cycles reagent kit (v1.5) in the Keck Genomics Platform (KGP) Core facility at USC.

The post-alignment statistics and quality measures of these libraries have been summarized previously ^[17]^.

### Variant calling

The sequencing reads are assessed and analyzed using a validated pipeline based on GATK v4.2 best practices from the USC Keck Genomics Platform (KGP) in-house pipeline, as described previously ^[1]^, and the DRAGEN Somatic v 4.2.4 pipeline (Illumina Inc.). PCR duplicate reads were identified and marked using Picard (MarkDuplicates) as part of the standard GATK preprocessing pipeline. These marked duplicates were excluded from downstream variant calling steps to prevent artificial inflation of variant support. Two germline callers, HaplotypeCaller and FreeBayes (v1.2.0), and two somatic variant callers, Mutect2 and Strelka (v2.9.0), are used in the KGP pipeline. Somatic variant calls in the crypt samples are made with the matched bulk tissue sample from the same individual as the control. The shared somatic variant calls from Mutect2 and Strelka (2Caller) through the KGP pipeline and the DRAGEN pipeline have a high inter-caller concordance with 93.78% (s.d. 2.79%) 2Caller calls present in DRAGEN and 80.77% (s.d. 4.92%) DRAGEN calls present in 2Caller. The variant calls from 2Caller are used for all downstream analyses to increase variant call confidence. The variant calls from Illumina’s DRAGEN somatic pipeline are only used for internal validation of the analysis.

### Simple deletion analysis

Deletions can derive from 1) strand slippage due to misalignment of repetitive sequences, such as homopolymer tracts or tandem repeats, during DNA replication or DNA breakage repair, leading to repeat contraction; 2) DNA breakage repair in regions of unique DNA sequence utilizing alignment of short homologous nucleotides; 3) DNA breakage in regions of unique DNA sequence repaired without any homologous sequence involved. The 2Caller variants are called by both Mutect2 and Strelka using a matched bulk tissue control from each individual. Only the common pass calls from Mutect2 and Strelka are included in the deletion variant list to ensure the high quality of the variants included in the analysis. A total of 24,181 autosomal simple deletion variants identified in 106 colon crypts from 21 individuals by the 2Caller are analyzed (Fig. 1). The duplicate variant calls present in more than a single individual (apparent inter-sharing) are removed and all the duplicate variant calls except one for each variant present in more than one crypt of a single individual (intra-sharing) are removed. After removing duplicate variants, the 23,145 variants are subjected to the hidden inter-sharing identification process as described previously with the modification of matching variant position in 2Callers to (position minus 1) in FreeBayes VCFs ^[1]^. The final 12,857 unique deletion variants are further analyzed.

**Figure 1.**
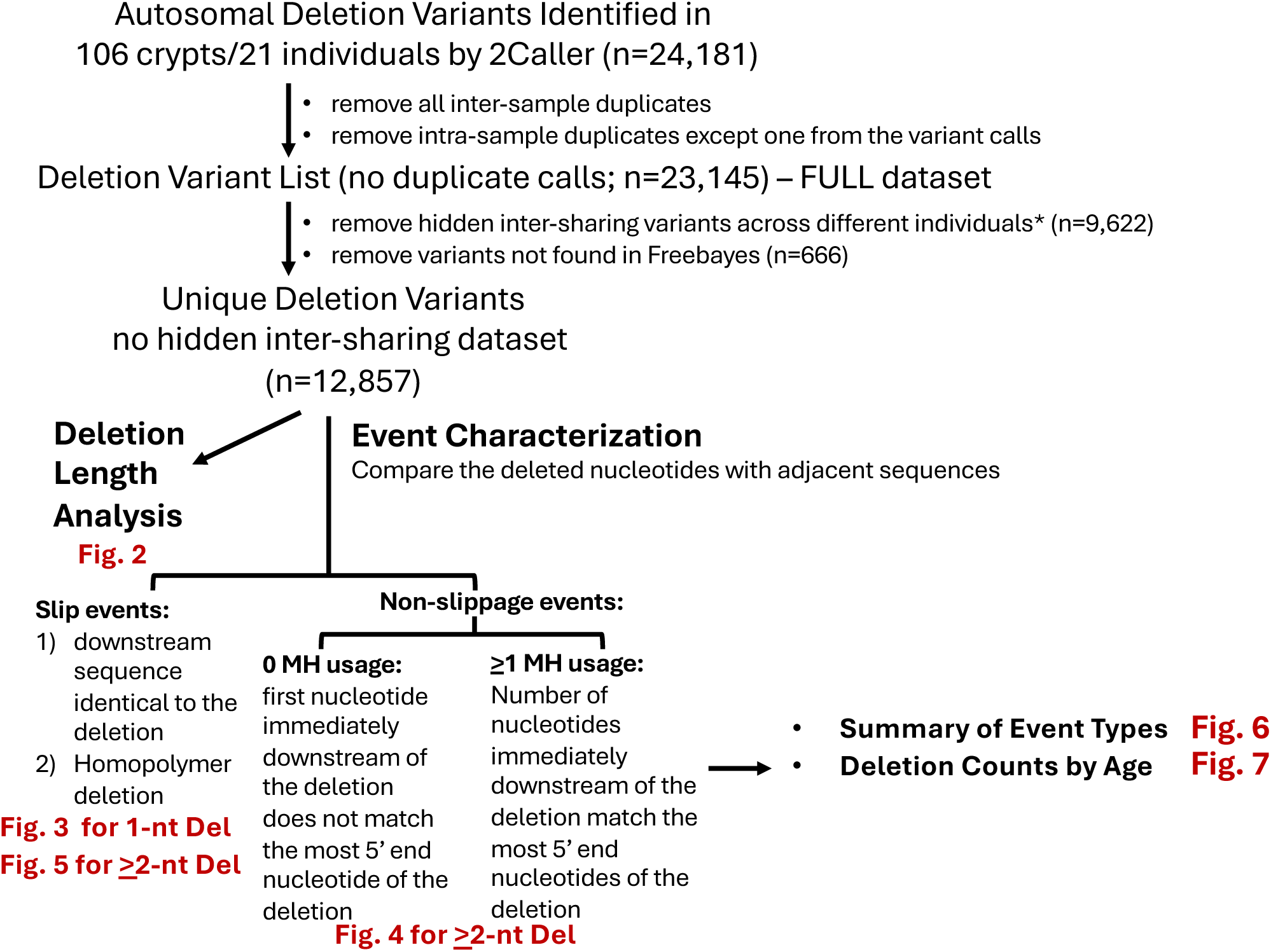
Deletion Analysis Flow Chart. The workflow for the analysis is summarized with the relevant Figures for the results indicated in **red**. See Methods and Materials section for details of the analysis.

Sequences immediately downstream of the deletions are compared with the deleted nucleotides to determine whether a deletion is a slip event or a non-slippage event. Indels are left-aligned according to VCF conventions, consequently the nucleotide immediately upstream of the deletion would never match the first nucleotide of the deletion. Therefore, only the sequence immediately downstream of the deletion is used for the slippage and microhomology analyses. Unlike the insertions, which can contain random and unknown sequences, the identity of deleted nucleotides can be clearly determined despite the exact location being ambiguous for variants in repetitive regions. The characterization of polymerase slippage or microhomology usage at each deletion junction is determined using a script (https://github.com/eddieloh-usc/SimpleDeletionAnalysis) developed based on the algorithms as follows; 1) Mcall step: The nucleotides along the deleted sequence are sequentially compared against the nucleotides (of length equal to the deleted sequence) immediately following the deletion position. When the sequence of the two nucleotides compared are identical, it is marked as “M” for that deleted nucleotide. When the sequence of the two nucleotides compared are different, the position of the deleted nucleotide is marked as “X.” The comparison stops when the first “X” appears in the string. 2) Scall step: a deletion is marked as “NA” and this step completes when nucleotides in the deletion are not all identical (i.e. not a homopolymer). Otherwise, similar to the Mcall step, the nucleotides along each deleted sequence are sequentially compared against the nucleotides (of length equal to the deleted sequence) immediately following the deletion position.

When the sequence of the two nucleotides compared are identical, it is marked as “S” for that deleted nucleotide. When the sequence of the two nucleotides compared are different, the position of the deleted nucleotide is marked as “X.” The comparison stops when the first “X” appears in the string. 3) The variants with any “S” in the Scall are counted as slip events of the homopolymer tract category. When the number of Ms in the Mcall is equal to the length of the deletion and Scall is NA, the variant is defined as a slip event of the tandem repeat category. The junctional microhomology is determined by the number of “M” in the Mcall when the number of Ms in the Mcall is smaller than the length of the deletion and the Scall is “NA” or “X”. The Mcalls and Scalls are listed in Supplementary Table S1 and summarized by the event type designated as slip and the number of microhomology usage (Supplementary Table S2).

## RESULTS

### A majority of simple deletions are 1-nucleotide deletions and >87% of the deletions are ≤5-nucleotide in length

Among the total of 12,857 unique deletion events (no variant inter-sharing, apparent or hidden), 8,593 (66.84%) variants are single nucleotide deletions (Fig. 2 and Supplementary Table S2). Deletion lengths up to 5-nucleotide account for nearly 87.74% of all simple deletions (Fig. 2A, Cumulative % of Total). The event count clearly decreases with increasing deletion length (Fig. 2A and 2B). The longest deletions observed are two variants with a 48-nucleotide deletion (Fig. 2A and 2B), likely due to the detection limit at 50 bases for deletions by both Mutect2 and Strelka. Less than 1% of the deletions are >25-nucleotide in length (Fig. 2A, Cumulative % of total = 99.03% for ≤25-nucleotide deletions). This finding indicates that large somatic deletions are uncommon in the human genome of normal (non-neoplastic) cells.

**Figure 2.**
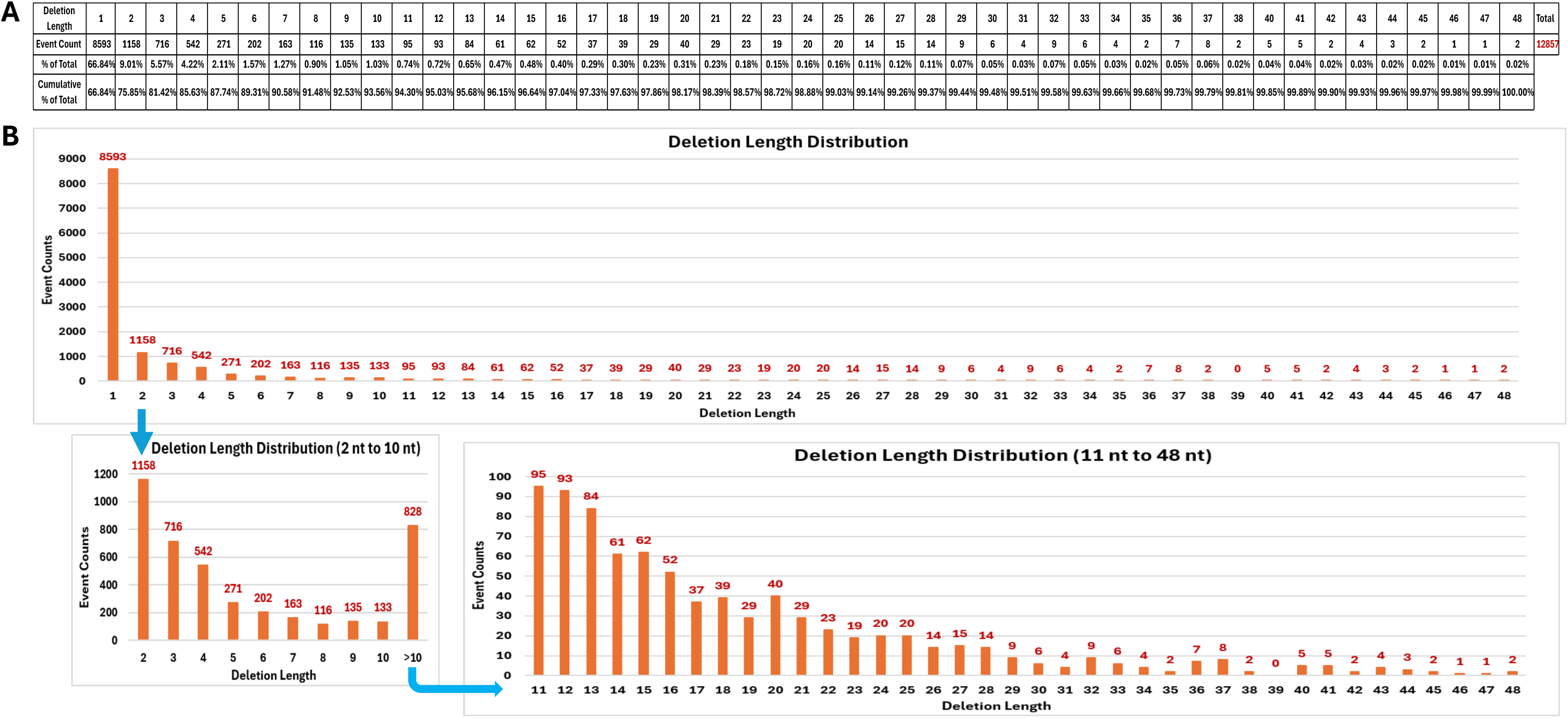
Deletion Length Analysis. Length of each deletion is counted and tabulated in A) with presentation of each deletion length calculated as percentage of the total. The distribution of the deletion events is plotted in B) with the number of events for each deletion length above the histogram.

Strand slippage events can be identified by examining sequences immediately downstream of the deletions for the deleted nucleotides. When the nucleotide followed immediately downstream of the deleted nucleotide is identical to the deleted nucleotide, the strand slippage event of the homopolymer tract length of two is the most likely mechanism for the deletion. Among the 8,593 1-nucleotide deletions, 6,353 (73.93%) have an identical nucleotide immediately downstream of the deleted nucleotide, indicating potential strand slippage in homopolymer tract (Fig. 3A, numbers in green).

**Figure 3.**
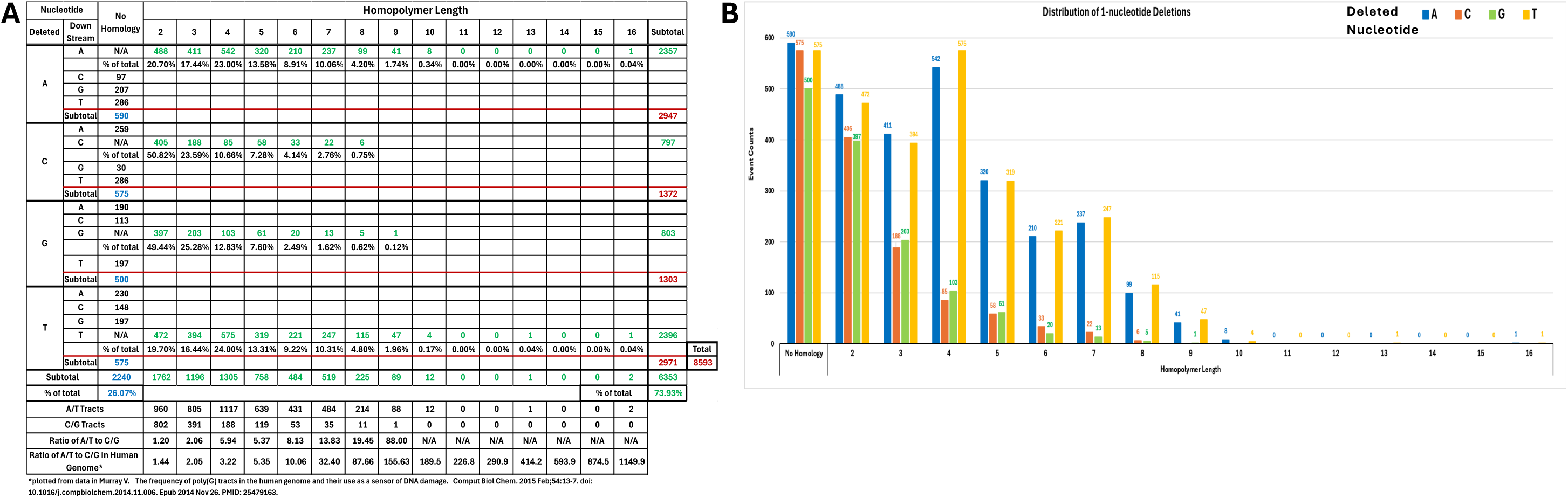
Summary of 1-nucleotide Deletions by Event Types. A) Count summary. Deletions of 1-nucleotide with one or more identical nucleotides immediately downstream of the deleted nucleotide is considered in the homopolymer tract. All other deletions of 1-nucleotide are grouped as no homology. The event count is grouped by nucleotide content of the deleted nucleotide and the immediate downstream nucleotide. Deletions in the homopolymer tract is counted by tract length including the deleted nucleotide. The counts and subtotal of deletions with no homology are in **blue**, the counts and subtotal of deletions in the homopolymer tracts are in **green**. The total of each deleted nucleotide and the total of all the 1-nucleotide deletions are in **red**. B) The distribution of 1-nucleotide deletions according to the identity of the deleted nucleotide are plotted against homopolymer length.

The remaining 2,240 1-nucleotide deletions (26.07%) do not share homology with the nucleotide immediately downstream and are considered non-slip deletions (Fig. 3A, number in blue). Examination of the nucleotide sequence further downstream of these events reveals that nearly all the slip deletion events occur in homopolymer tracts shorter than 10 and occur more often at A or T (A/T) tracts than C or G (C/G) tracts (Fig. 3A and 3B). It is notable that prevalence of the 1-nucleotide deletions in A/T tracts becomes much higher than in C/G tracts when homopolymer length is ≥3. This observation is expected as homopolymers are rarely longer than 10 nucleotides in the human genome. The ratio of deletions in A/T to C/G tracts observed in this study is similar to the homopolymer tract ratio in the human genome for tract length <5-nucleotide. However, deletions of >5-nucleotide in the A/T tracts are less frequently observed than expected based on A/T homopolymer tract representation in the human genome ^[21]^. Among the non-slippage deletions, the content of the deleted nucleotide makes no appreciable difference (Fig. 3A numbers in blue & 3B “No Homology” column). However, it is impossible to distinguish whether the event is due to strand slippage or a randomly occurring repair event at a location with two identical adjacent nucleotides. Since the mechanism cannot be clearly inferred for the single nucleotide deletions, further analysis presented below is restricted to deletions ≥2 nucleotides.

### Microhomology ≥5-nucleotide is uncommon in the ≥2-nucleotide deletions

When the deletion is ≥2-nucleotide, homopolymer tracts and tandem repeats can lead to strand slippage by misalignment during DNA replication or DNA damage repair and result in deletions. As described in the Methods, a homopolymer tract is signified by the presence of ≥1 consecutive nucleotides identical to the deleted nucleotides immediately downstream from the deletion. Tandem repeat deletion events are defined by the presence of one or more units of contiguous nucleotides immediately downstream of the deletion that are identical to the entire deleted sequence, also as described in the Methods. Deletion events without the slippage signature are DNA damage repair events that can be with or without microhomology (MH) usage at the junction of repair depending on the repair pathway. Microhomology is identified by comparing the deleted nucleotides with the nucleotides immediately downstream for maximum homology, as described in the Methods.

Among the 4,264 ≥2-nucleotide deletions (Fig. 4A, number in red), 1,157 (27.13%, Fig. 4A, numbers in blue) are slip events. Also present, 1,040 (24.39%, Fig. 4A, numbers in orange) do not have MH immediately downstream from the deletion, and these are designated 0 MH (defined as cases where the nucleotide immediately downstream from the deletion is not identical to the first nucleotide of the deletion).

**Figure 4.**
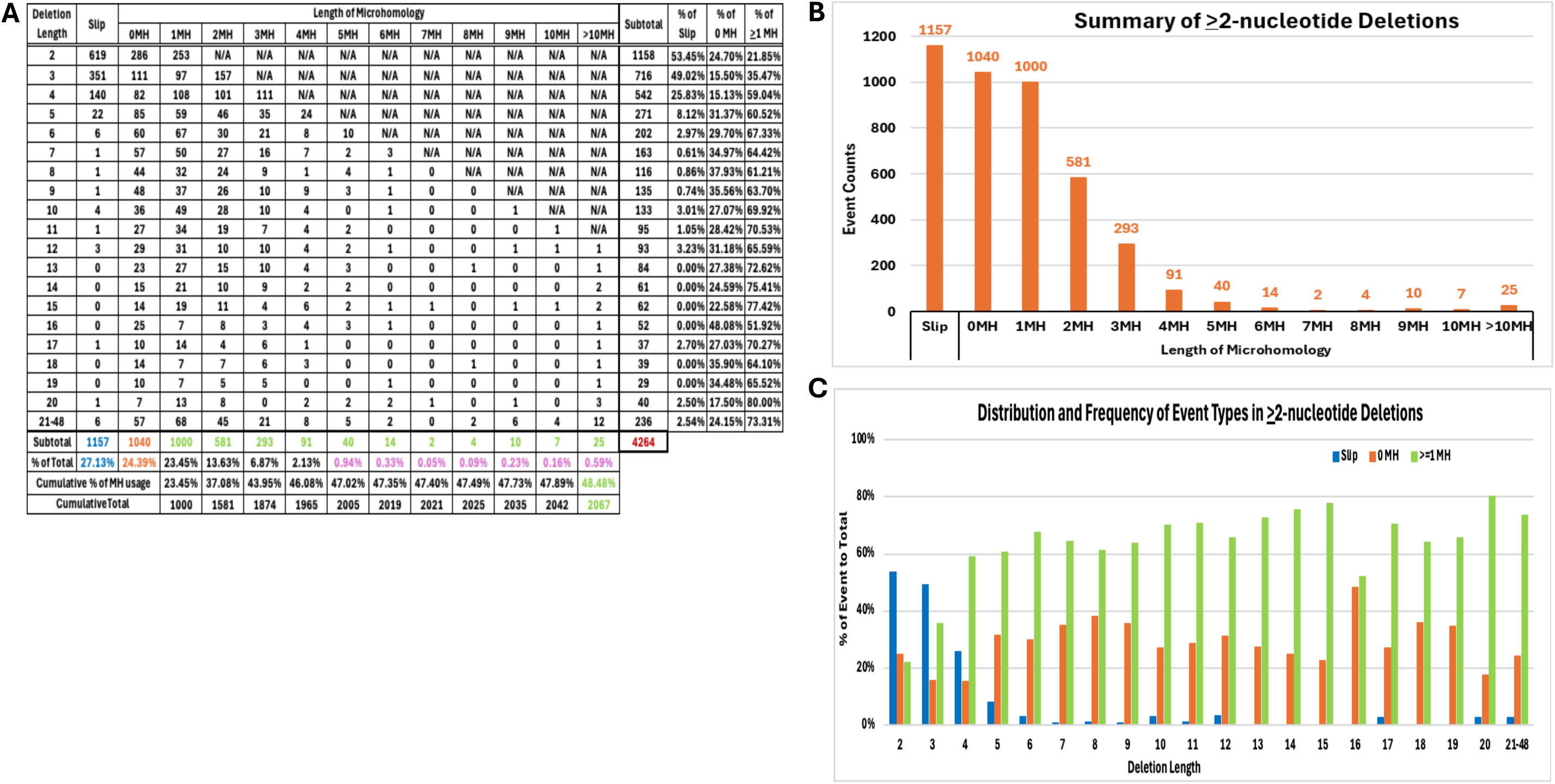
Event Counts by Sequence Features in the ≥2-nucleotide Deletion Junctions. A) Event counts of deletions by features of strand slippage and microhomology immediately downstream of the deletion. The total count and frequency of slip events are in **blue**, those of the events with no homology immediately downstream of the deletion are in **orange**. The total count of the events with ≥1-nucleotide microhomology are in **green,** and the frequency of events with ≥5-nucleotide microhomology are in **pink**. The total of number of deletions included in the analysis is in **red**. The complete table of microhomology frequency calculation is presented in Supplementary Table S3. B) Graphical illustration of deletions by event types. C) Graphical illustration of deletion event types and their frequencies by deletion length.

Most abundant, 2,067 (48.48%, Fig. 4A, numbers in green) showed ≥1 nucleotide MH at the junction of the deletion. Among these deletions with any MH (2,067 events, 48.48% of the total), 1,965 of the events have <5-nucleotide MH, and each MH length ≥5-nucleotide only accounts for <1% of the total events (Fig. 4A, numbers in pink). These findings indicate that MH >5-nucleotide is rarely present at deletion junctions.

When deletion events of each length are examined, it is clear that the presence of slip events decreases as the length of deletion increases. In contrast to the 2- and 3-nucleotide deletions, among which ∼50% of the events are slip events, slip events only account for ≥5% of the ≥5-nucleotide deletions (Fig. 4A and Fig. 4C, blue histogram).

Among the ≥4-nucleotide deletions, >50% of the events have MH at the deletion junctions (Fig. 4A and Fig. 4C, green histogram). The presence of slip events and events with MH at the deletion junction reverses in predominance when the deletion length is ≥4-nucleotide. Interestingly, the presence of non-slip events with 0 MH does not change drastically across the deletion length (Fig. 4A and Fig. 4C, orange histogram). These findings indicate that deletions ≥4-nucleotide are less likely due to DNA replication errors and more likely to be the result of DNA repair events.

### A majority of the ≥2-nucleotide slip deletions are in the tandem repeats

The 1,157 slip deletions include 273 (23.60%, Fig. 5A and Fig. 6) deletions within homopolymer tracts and 884 (76.40%, Fig. 5B and Fig. 6) deletions involving tandem repeats. The longest homopolymer tract deletion observed in the 273 events is 18 nucleotides in length, and homopolymer tract deletions >7-nucleotide are uncommon (Fig. 5A). The A/T homopolymer tracts are observed much more often and of longer length than the C/G homopolymer tracts among these 273 deletions (Fig. 5A). In the human genome, the ratio of A/T to C/G homopolymer tracts increases dramatically as the homopolymer tracts become longer (Fig. 5A) ^[21]^. The ratio of deletions in the A/T to C/G homopolymer tracts we observed in this study is ≥2-fold higher than the ratio of their respective presence in the human genome for homopolymer tract lengths of 3 and 4. However, the ratio of deletions of A/T to C/G homopolymers is closer to the presence of corresponding tracts in the human genome for 5- and 6-nucleotide tract lengths (Fig. 5A).

**Figure 5.**
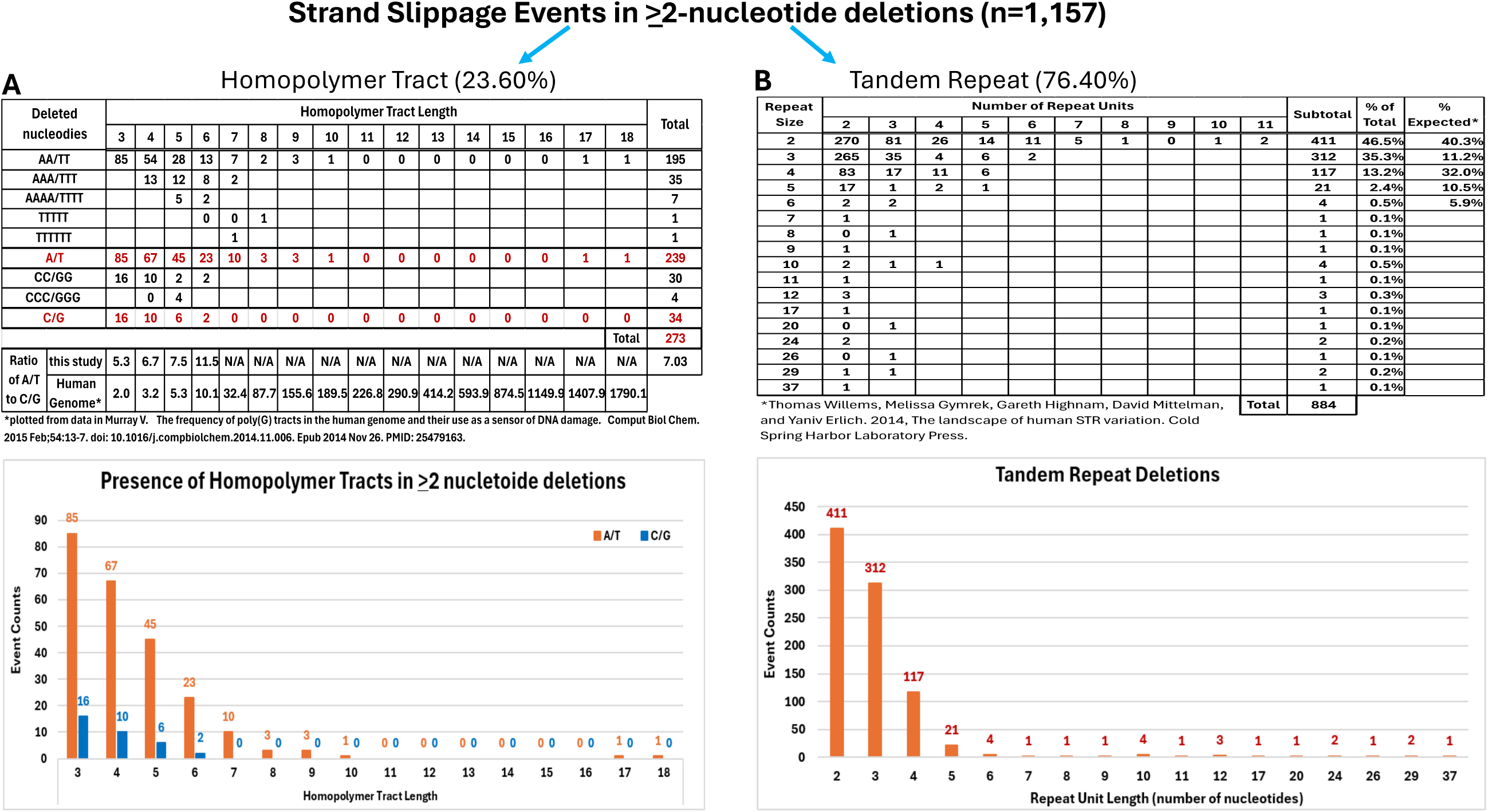
Strand Slippage Events in ≥2-nucleotide Deletions. A) Event counts of deletions occurring in homopolymer tracts by the deleted nucleotides and the ratio of A/T to C/G tracts by tract length are summarized in the top panel. The event counts in the A/T and C/G tracts are plotted against the homopolymer tract length in the bottom panel. B) Event counts of deletions occurring in the tandem repeats are summarized in the top panel by repeat unit length and the number of repeat units. The total event counts for each repeat length are plotted in the graph in the bottom panel.

**Figure 6.**
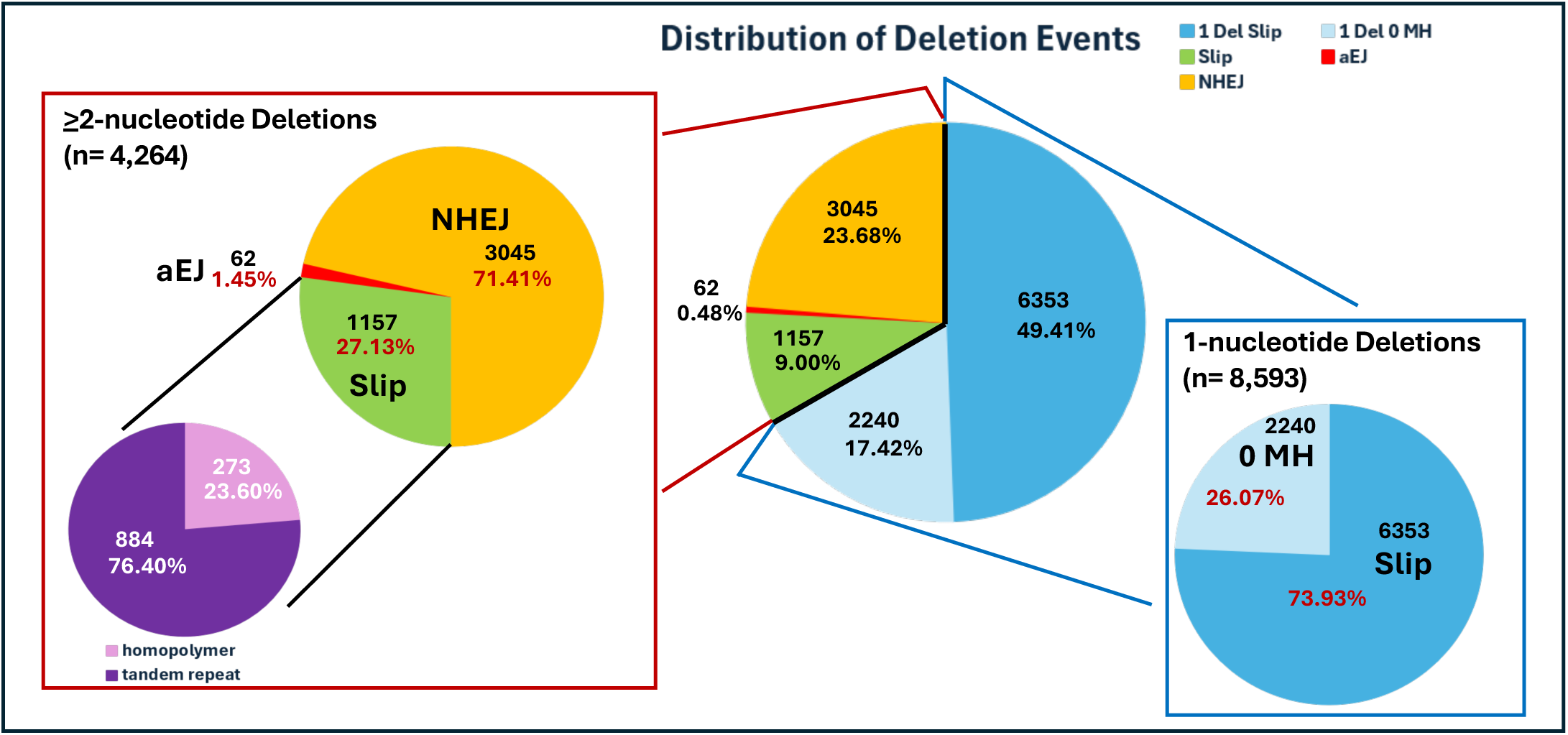
Distribution of Deletion Events by Mechanisms. For 1-nucleotide deletion events: ‘Slip’ means that the nucleotide immediately downstream is identical to the deleted nucleotide. ‘0 MH’ means that the nucleotide immediately downstream is different than the deleted nucleotide (0 homology). For ≥2-nucleotide deletion events: “slip” means that there is perfect match of sequences immediately downstream to the deleted nucleotides or means homopolymer deletions with at least one identical nucleotide immediately downstream. aEJ: deletion events using microhomology of length 6 to 31 nucleotides. NHEJ: deletion events using microhomology of 0 to 5 nucleotides in length. Among the ≥2-nucleotide deletions, four events can potentially be stem-loop excision events with a 5-nucleotide stem: 3 events are included in the NHEJ section of the pie graph and one is included in the aEJ section of the pie chart, all using microhomology.

Dinucleotide (n = 411, 46.5%), trinucleotide (n = 312, 35.3%), and tetranucleotide (n = 117, 13.2%) repeats account for 95% of the 884 deletions involving tandem repeats (Fig. 5B). The presence of the di-, tri-, and tetra-nucleotide tandem repeats in the deletion events are dissimilar to their presence in the human genome (40.3%, 11.2%, and 32.0%, respectively, among the 2- to 6-nucleotide repeats), especially for trinucleotide repeats (Fig. 5B) ^[22]^.

### A majority of the ≥2-nucleotide non-slippage deletions are NHEJ events

The ≥2-nucleotide non-slippage deletions (n = 3,107) include events without any MH at the deletion junction and events with one or more homologous nucleotides matching the deletion. While NHEJ does not rely on extensive DNA end microhomology to join broken DNA ends, NHEJ can use 0- to 5-nucleotide of microhomology for end joining ^[5]^. In contrast, microhomology >5-nucleotide and up to ∼30-nucleotide are more commonly associated with aEJ DNA repair ^[6]^. We classified our set of 3,107 non-slippage deletions into NHEJ events with 0- to 5-nucleotide of MH (n = 3,045) and aEJ events with >6-nucleotide (n = 62) of MH between the deleted nucleotides and the nucleotides immediately downstream of the deletion (Fig. 4A and Fig. 6). The rationale for this classification is presented in the Supplementary Information and the Supplementary Fig. S1 within. Two points are clear. First, NHEJ (71.41%) occurs at a much higher frequency than aEJ (1.45%) for DNA repair in the colon epithelium, consistent with the low expression of pol theta (Fig. 6). MH >5-nucleotide is rarely observed at the deletion junctions in our study, also consistent with aEJ events being uncommon in colon crypts. Second, DNA damage events (71.41% + 1.45%) occur at a higher frequency than DNA replication strand slippage (27.13%) among ≥2-nucleotide deletions.

DNA sequence with palindromic or other distinctive sequence characteristics may also lead to deletions, unrelated to NHEJ and aEJ. For example, when a local intra-strand palindromic DNA pairing allows a stable stem-and-loop conformation to form within each strand of the duplex, the DNA can be excised at the base of the stem leading to a deletion spanning the entire stem-and-loop. Among the 3,107 non-slippage deletions, the largest potential stem-and-loop size is a 5-nucleotide stem with the 5-nucleotide on the 5’-end being the reverse complement of the 5-nucleotide on the 3’-end. But we observed only three of the NHEJ events (Supplementary Table S1, rows 1399, 3315, and 3670 marked by yellow highlight) and a single aEJ event (row 11215 in Supplementary Table S1, marked by yellow highlight) that might potentially form a stem-and-loop with a 5-nucleotide stem. This observation indicates that stable stem-loop formation does not naturally lead to deletion, based on the low frequency of this DNA repair signature in the human genome of normal somatic cells.

### Deletions of ≥2-nucleotide accumulate with age

We learned in our previous analysis of accumulation of SNVs with age that clonality of the crypt can affect the variant allele frequency and mutation counts ^[18, 23]^. The rate of accumulation for non-slippage and slip deletions may be different since they are caused by different factors. Also, chemotherapy or chemo/radiation combination treatment affects DNA metabolism and can cause both non-specific and specific DNA damage. Therefore, the rate of deletion accumulation is calculated for each type of deletion in different groups of the study individuals and crypt clonality, as in our previous analysis of SNVs ^[23]^. Similar to autosomal SNVs and the complex indels ^[20, 23]^, ≥2-nucleotide simple deletions also accumulate linearly with age in 1SC colon crypts from individuals without chemotherapy or chemo/radiation combination treatment (Fig. 7). It is clear that chemo/radiation combination treatment affects non-slippage deletions much more than slip deletions as indicated by the very poor R^2^ for non-slippage deletions when individuals with treatment are included (compare Fig. 7A and 7B). Excluding individuals with treatment clearly increases the R^2^ (compare top and bottom panels in Fig. 7A, 7B, 7C, and 7D) for both types of deletions. The linear accumulation of deletions with age is observed in the ≥2-nucleotide non-slippage deletions at a rate of 0.34 per year with a R^2^ = 0.9057 and in slip deletions at a rate of 0.23 per year with a R^2^ = 0.9029 when all 70 crypts from individuals with no treatment are included in the analysis (Fig. 7A and 7B, bottom panels). Similar linear accumulation of non-slippage deletions at a rate of 0.3574 per year with a R^2^ = 0.9201 and slip deletions at a rate of 0.2234 per year with a R^2^ = 0.9476 is observed when only the 36 monoclonal crypts from individuals without treatment are included in the analysis (Fig. 7C and 7D, bottom panels). Unlike the SNVs, clonality of the colon crypts does not seem to impact the deletion counts appreciably, as indicated by the minimal increase of R^2^ when only the monoclonal crypts are included in the analysis. This may potentially be due to the very low rate of simple deletion accumulation at ∼0.22 to 0.36 per year compared with that of SNVs at ∼45 per year. It is noteworthy that the slope of the linear regression model, which reflects the rate of mutation per year, is consistently lower for the slip deletions (∼0.22 per year) than the non-slippage deletions (∼0.34 per year).

**Figure 7.**
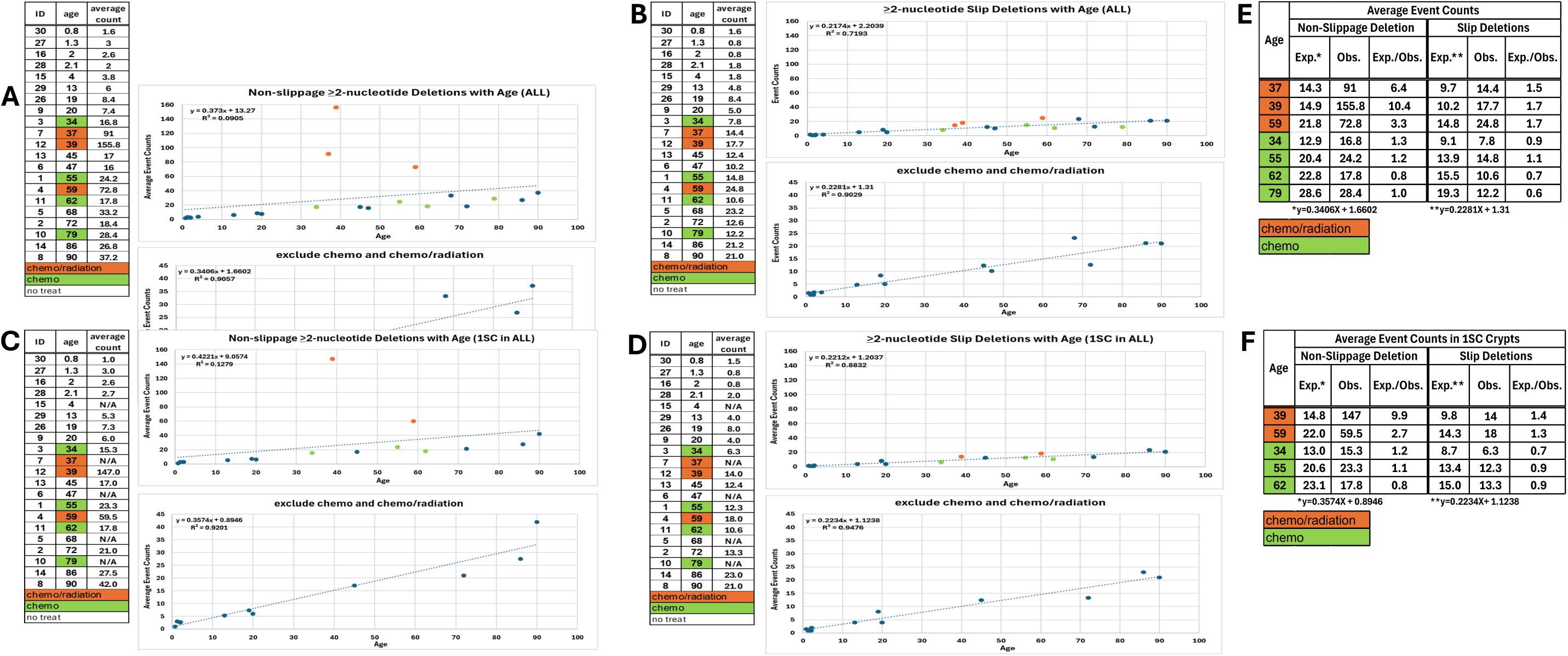
Average ≥2-nucleotide Deletion Counts Increase with Age. A) Top panel: average counts of ≥2-nucleotide non-slippage deletions in all colon crypts studied from each of the 21 individuals (106 crypts total) plotted against age. Bottom panel: replotting data in the top panel after removing the 7 individuals treated with either chemotherapy or a chemo/radiation combination treatment (70 crypts total). B) Top panel: average counts of > 2-nucleotide slip deletions in all colon crypts studied from each of the 21 individuals plotted against age. Bottom panel: replotting data in the top panel after removing individuals with treatment. C) Top panel: average counts of ≥2-nucleotide non-slippage deletions in 1SC colon crypts studied from each of the 16 individuals with any 1SC crypts (52 crypts total) plotted against age. Bottom panel: replotting data in the top panel after removing the four individuals with treatment (36 crypts total). D) Top panel: average counts of > 2-nucleotide slip deletion in 1SC colon crypts plotted against age. Bottom panel: replotting data in the top panel after removing the four individuals with treatment. In A) to D), the **green dots** designate individuals with chemotherapy; the **orange dots** designate individuals with chemo/radiation combination treatment; and the **blue dots** designate individuals with no treatment. A linear trend line is displayed with the equation and the R-square value in all plots. E) and F) Increase of deletions with treatment are measured by comparing the observed even count (Obs.) with expected event count (Exp., “y” in the equation below each type of deletion) derived by using the age of the individual as “X” in the equation. The equations are from the linear regression model of the respective plots in A) to D).

### Radiation treatment increases non-slippage deletions

As mentioned above, the slip deletions are mostly due to strand slippage during DNA replication without DNA breakage and the non-slippage deletions are due to DNA breakage repaired predominately by NHEJ with few aEJ events. We observed a striking and interesting increase in non-slippage deletions in the individuals treated with chemo/radiation (Fig. 7A and 7C, orange dots in the top panels), whereas no dramatic increase is observed in the slip deletions (Fig. 7B and 7D, orange dots in the top panels). The expected deletion counts for each individual can be calculated based on the age of an individual and the slope (rate of deletion accumulation) and the intercept from the linear regression model. Deviation of the observed deletion event count from the expected deletion count can be calculated as the fold difference (the observed count divided by the expected count). The three individuals with chemo/radiation combination treatment have 3.3- to 10.4-fold more non-slippage deletions than expected when all crypts, regardless of their clonality, from each individual are included; in contrast, there is only a ∼1.5-fold increase of slip deletions than expected (Fig. 7E). Similarly, there are 2.7- and 9.9-fold more non-slippage deletions than expected when only monoclonal crypts are included in the calculation (the 39-year-old individual does not have any 1SC crypt) as presented in Fig. 7F. Again, slip deletion only shows a very small difference between the expected and observed deletion count in the monoclonal crypts from the two individuals with the chemo/radiation combination treatment (Fig. 7F). Interestingly, individuals treated with only chemotherapy showed no appreciable difference between the expected and observed deletion count in both deletion types, regardless of clonality (Fig. 7E and 7F). It is well established that radiation causes DNA damage ^[3, 5, 24]^. These data are consistent with radiation being the principal contributor to the increased non-slippage deletions in the chemo/radiation-treated individuals, because a comparable increase was not observed in individuals treated with chemotherapy alone.

## Discussion

There are several conclusions from the analysis of simple deletions in human colon crypts. First, ∼67% of the simple deletions are 1-nucleotide deletions and ∼74% of these 1-nucleotide deletions are slip deletions. Second, among the ≥2-nucleotide deletions (i.e. the remaining ∼33% of the simple deletions), MH of >5-nucleotide is uncommon. Third, deletions >4-nucleotide are mostly the result of DNA breakage repair by NHEJ with a small number by aEJ, while the shorter deletions are mostly due to strand slippage. Fourth, ≥2-nucleotide deletions accumulate with age in the colon crypt stem cells. Fifth, radiation treatment causes DNA breakage and leads to mainly non-slippage deletions, which can potentially be measured and distinguished from a well-established baseline.

### Single Nucleotide Deletions

Simple deletions of only 1-nucleotide can arise due to either strand slippage or a wide variety of single-strand damage events that are subject to base excision repair (BER). Single nucleotide deletions in the homopolymer tracts account for 49.41% of the simple deletions in our study (Fig. 6 which is 73.93% of the 1-nucleotide deletion events), clearly indicating that slippage is a dominant cause of 1-nucleotide deletions.

Strand slippage is most commonly associated with DNA synthesis, particularly during cell replication. The presence of C/G homopolymer tracts in the human genome decreases relative to A/T tracts as the tract length increases ^[21]^. Among the 6,353 deletions at homopolymer tracts observed in our study (Fig. 3), the A/T to C/G ratio does not rise nearly as much as the representation of these two tracts in the human genome when the tract length is 6- to 8-nucleotide. It has been documented that A-tracts of length 4- to 9-nucleotide have more stable bends in duplex DNA than other tract lengths ^[25, 26]^. The stability of A/T homopolymer tract bends may be the underlying reason for the much lower ratio of A/T to C/G tracts observed for deletions in tract lengths of 6- to 8-nucleotide.

The other 26.07% of the 1-nucleotide deletions (17.4% of the total deletions) are not in any homopolymer tracts; therefore, there is no microhomology (MH) immediately downstream (Fig. 6). These 0 MH deletions occur at similar frequencies for A, C, G, and T nucleotides. This equal nucleotide representation may arise from a combination of factors, including random probability. Oxidative damage giving rise to 8-oxo-G is one factor favoring 1-nucleotide deletions during BER at G/C sites ^[27]^. A bacterial toxin, colibactin, described in normal human colon crypts of colon cancer patients favors deletion of nucleotide A ^[28]^. Additional studies will be needed to understand the basis for the nucleotides prone to single base deletion.

### ≥2-nucleotide Deletions

Slippage in homopolymeric or tandem repeat sequences accounts for 9% of the total simple deletions and 27.13% of the ≥2-nucleotide deletions (Fig. 6). The non-slippage deletions include deletions with no homologous nucleotide immediately downstream of the deleted nucleotide (0 MH) and deletions with microhomology immediately downstream (>1MH). While slip deletions are the predominant events in the 2- and 3-nucleotide deletions as in the 1-nucleotide deletions, the majority of the >4-nucleotide deletions have at least one nucleotide of microhomology (MH) immediately downstream from the deleted nucleotide, most consistent with an NHEJ mechanism (Fig. 4).

Similar to the 1-nucleotide deletions, the frequency of deletions in the homopolymer tracts decreases as tract length increases. There are no deletions observed in the C/G homopolymer tracts of >6-nucleotide in length; therefore, the ratio of deletion events in A/T to C/G tracts of >6-nucleotide cannot be compared with that of 1-nucleotide deletions. The majority of the ≥2-nucleotide slip deletions are in the tandem repeats (76.40%) with ∼95% of the deletions within di-, tri-, and tetra-nucleotide repeat tracts. The number of events decreases with the unit length of the repeat as well as the number of repeat units. Interestingly, the presence of deletion in the di-, tri-, and tetra-nucleotide repeats is discordant with their abundance in the human genome.

These observations support that DNA stability drives slip deletion of tandem repeats and the abundance of the repeats in the genome only plays a minor role.

Most of the ≥2-nucleotide deletions are non-slippage events (72.87%) and primarily have 0 to 2 MH (Fig. 4A, 24.39% plus 37.08%) immediately downstream of the deleted nucleotides. The mechanism for these predominant events is most consistent with NHEJ. NHEJ can include insertions and base changes beyond simple deletions, namely complex indels. However, most of the variant callers are not designed to identify complex indels. We developed an algorithm to extract potential complex indels from FreeBayes and have reported the findings previously ^[19]^. It is noteworthy that there are 3,107 non-slip ≥2-nucleotide simple deletions reported here and 385 complex indels identified having a specific NHEJ repair signature from the 21 individuals studied^[19]^. It appears that simple deletions occur nearly 8-fold more often than complex indels.

In previous biochemical NHEJ studies with purified proteins including pol mu, pol lambda, and TdT, complex indel repair is the predominate outcome. However, if TdT is absent from the assay, simple deletions and complex indels are observed at similar frequencies ^[29, 30]^. In naturally-occurring V(D)J recombination events in human cells, where TdT is commonly present, complex indels are observed more often than simple deletions. In transfection V(D)J recombination assays using cells without TdT, most of the DNA double strand breaks result in simple deletions ^[31]^. It is important to note that the active polymerase concentration in the biochemical assays is likely different from what is in colon crypt cells, which do not express TdT, but express pol mu and lambda.

Our complex indel algorithm focused only on deletion with nucleotide insertions shorter than the length of the deletion; therefore, the 385 complex indel events are obviously an underestimate of the NHEJ activity, even though this would not entirely account for the 8-fold difference in simple deletion and complex indel events we identified from this dataset. However, the high ratio of simple deletions to complex indels in the single colon crypts suggests that simple deletions are the most common double-strand break repair signature, parallel to what has been observed in the cellular assay of V(D)J recombination when TdT is not present ^[31, 32]^. It is possible that less complex damage does not require nucleotide addition for end repair as frequently as complex damage.

Naturally occurring DNA damage in normal cells in the human body (not cells in culture and not neoplastic cells) may generate less complex DNA features than the breakage generated by V(D)J recombination or ionizing radiation. It will be interesting to understand what features at DNA ends determine whether NHEJ repair invokes the error-prone pol X polymerases, specifically pol mu and lambda. The involvement of error-prone pol X polymerases may dictate the abundance of complex indels in other biological settings such as at V(D)J recombination junctions or in ionizing radiation repair ^[3–5, 33]^. In fact, the causes of naturally-arising DNA breaks in normal cells still need further investigation. Our single colon crypt study demonstrates the capability to identify somatic mutations in human stem cells comprehensively. Future studies of clonal cell populations from humans may allow the identification of natural causes of DNA breakage.

Among the ≥2-nucleotide non-slippage deletions, 24.39% and 23.45% of the events involve 0- or 1-nucleotide of MH, respectively. Furthermore, only <3% of the ≥2-nucleotide non-slippage deletions have >4-nucleotide of MH. NHEJ most often uses 0- or 1-nucleotide of MH and rarely uses more than 5-nucleotide of MH other than in repetitive sequences. The frequent occurrence of 0- and 1-nucleotide MH and the drop-off of MH length >5-nucleotide supports NHEJ involvement rather than aEJ (Fig. 4B and Supplementary Information) ^[6]^. This is consistent with the very low level of polymerase theta in normal human colon epithelium, in contrast to neoplastic colonic epithelium.

Like all neoplasms and rapidly dividing cells, polymerase theta level in colon carcinoma cells is higher than in normal cells; therefore, aEJ can occur more frequently in neoplasms ^[6]^. Also of note, we do not observe any clearly longer MH cases that would suggest single-strand annealing ^[3]^.

### Age-Related Change in the NHEJ and aEJ Events

When the average event count of ≥2-nucleotide non-slippage deletions (which are predominantly NHEJ events) from each individual is plotted against age, a linear trend of accumulation with age at a rate of 0.36 per year (slope in Fig. 7C, bottom panel) is apparent in the total of 36 monoclonal colon crypts from 11 individuals without any chemotherapy or a combination chemo/radiation treatment. The rate of simple deletion accumulation is 0.34 when based on all clonality crypts from 14 individuals with no treatment (70 crypts). These two calculations suggest that the clonality of the crypts may not affect the calculation dramatically. This rate of accumulation calculated previously for the complex indels, which are also predominantly NHEJ events, is 0.19 ^[1, 20]^. There are many fewer complex indels than simple deletions identified; therefore, the rate of NHEJ event accumulation is calculated with total event counts in all crypts from individuals with no treatment (70 crypts). Considering that the complex indels we identified previously are only a subset of all NHEJ events (a higher event count would lead to a higher mutation rate) and the difference in rate calculation, these two calculated results are very similar.

Although NHEJ events occur much less frequently than SNVs, NHEJ events have the potential for a larger impact when occurring in coding regions because they can change the reading frame and are more likely to cause truncation mutations. Even in non-coding regions harboring regulatory sequences such as promoters and enhancers, the simple deletions and complex indels of NHEJ have greater potential effects relative to SNVs. For this reason, both the simple and complex indels may be of substantial importance in their genomic impact on human aging.

### Radiation leads to a measurable increase of non-slippage deletions

A striking increase of non-slippage deletion events was observed in colon crypts from all three individuals with chemo/radiation combination treatment was observed (Fig. 7). This outcome appears to be primarily driven by the radiation treatment since very little change was observed in individuals with only the chemotherapy that was given in these cases. Importantly, radiation treatment does not have any appreciable effect on slip deletions. It is well established that radiation causes DNA damage ^[3, 5, 24]^. Our study presented here uses direct whole genome sequencing of single colon crypts without any ex vivo culture or DNA extraction and with bulk DNA control from the same tissue of each individual; therefore, naturally occurring somatic variants in the progenitor cell (colon crypt stem cell) that gives rise to the crypt lining can be identified. The analyses of these events naturally occurring in the human body provide clear support that the chemotherapy treatment here does not lead to increased DNA breakage; therefore, it does not increase non-slippage deletions. In contrast, the non-slippage deletions, which involve DNA breakage, reflect the effect of DNA damage by irradiation of cells within the human body. These findings also demonstrate that the effect of DNA damage by radiation can potentially be assessed by the increase of the non-slippage deletions. Potentially, studies may be designed to provide information on the threshold for DNA breakage induced cell death. Further studies are of high priority to confirm and expand our findings for clinical applications.

## ACKNOWLEDGEMENT

We would like to thank the Norris Comprehensive Cancer Center Translational Pathology Core for the sample collection. We would like to acknowledge the Norris Comprehensive Cancer Center Molecular Genomics Core and the Keck Genomics Platform at the University of Southern California for the sequencing work. This work is supported by funds from NIA R01 AG 067615, NIA R56 AG 067615, and the Catherine and Joseph Aresty Endowment to CLH. MRL was supported by NIGMS R35 118009.

## Supplemental Information

### Rationale for NHEJ and aEJ classification based on microhomology length

Most cellular data regarding inferred MH usage is derived from neoplastic cells or other rapidly dividing cells in culture. These rapidly dividing cells typically have higher pol theta expression than many normal human somatic cells. Particularly, the normal human colon epithelial crypt cells that we have studied here are known to have very low expression of pol theta. For these reasons, our data on MH presence in simple deletions naturally arising in single normal human colon crypts taken from the human body directly for WGS analysis without DNA extraction or WGA permits a view of end joining in its natural context without any extrinsic modulation.

In Supplementary Figure S1A, the event counts decreases from 0 MH to 6-nucleotide very quickly from 1041 to 14. In the bottom panels of Figure S1B, C, and D, a small peak around 9- to 11-nucleotide MH clearly arises among 7-nucleotide to 31-nucleotide MH. This rise in event counts among the >6-nucleotide MH is most consistent with aEJ. Considering that there are only two 7-nucleotide events and the rising of the small peak starts at 7-nucleotide, NHEJ is not likely to contribute to any deletions associated with 7-nucleotide MH. However, the 6-nucleotide MH is likely a mixture of NHEJ and aEJ. The low expression of pol theta in normal human colon epithelial crypt cells (non-neoplastic) is consistent with the much lower count of aEJ events observed in this study and is consistent with recent biochemical studies of pol theta mechanism ^[1, 2]^. Therefore, we choose to classify NHEJ events in this study as < 5-nucleotide MH and count MH length 6-nucleotides or longer as aEJ events for these deletions arising within the human body in normal colon crypt cells.

---

**Supplementary Figure S1.**
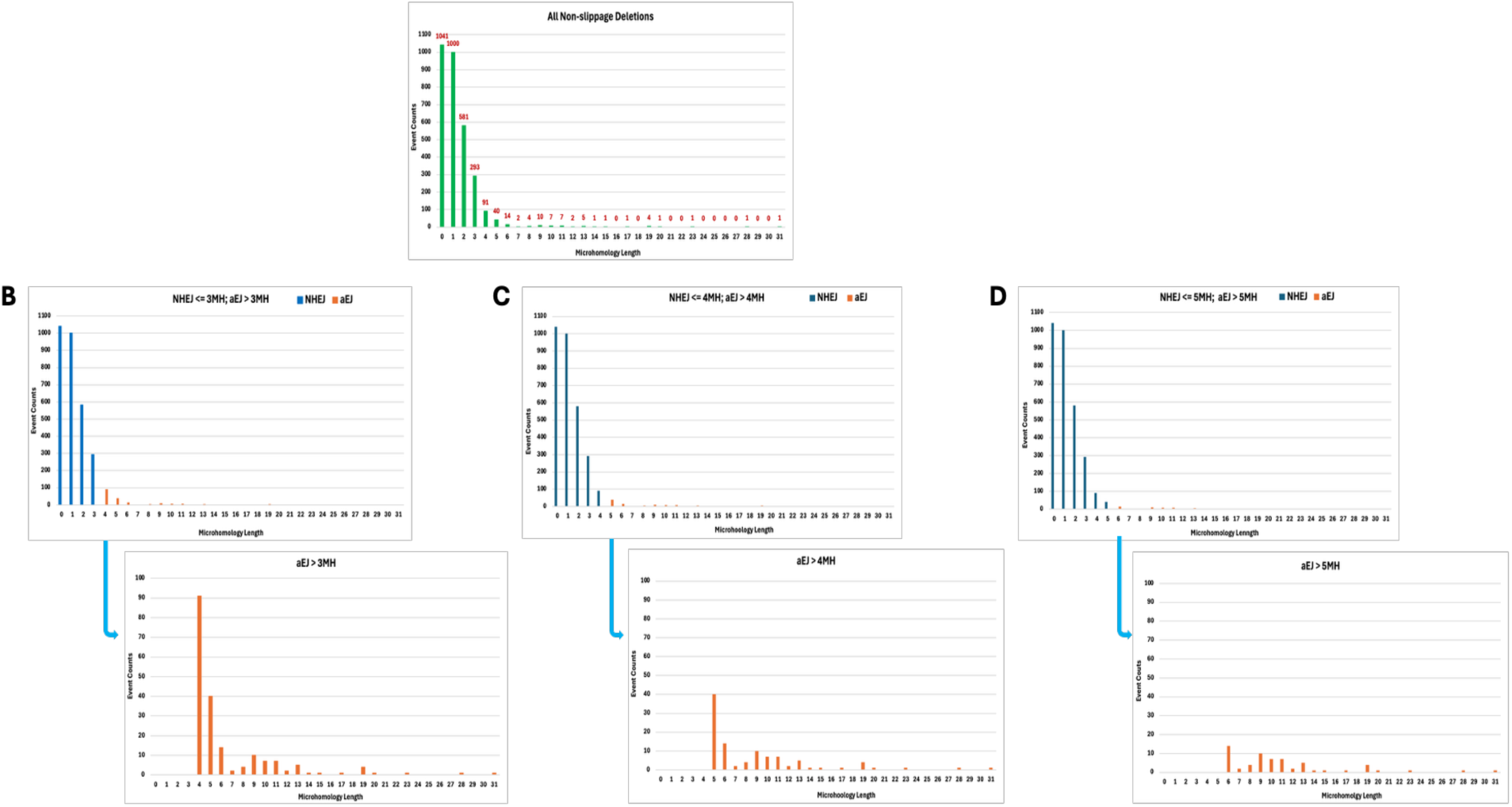
Rationale for NHEJ and aEJ classification based on microhomology length. A) The deletion event counts are plotted as a function of the microhomology (MH) length. The event counts of the non-slippage ≥2-nucleotide deletions decrease as microhomology (MH) length increases forthe total observed set of events. NHEJ and aEJ both use MH for end joining, and NHEJ is commonly observed with shorter MH length than aEJ. An overlapping zone of MH lengths can be associated with the two end joining processes, but ata lower frequency than the more favorable MH lengths for each process. To illustrate this point, the often debated MH cutoff length between 4- and 6-nucleotide are considered. The blue histogram bars are taken to be NHEJ events, and the orange bars are taken to be aEJ events. The aEJ-type events are infrequent in all cases and therefore their plots are expanded in the histogram below each combined plot in parts B, C, and D. B) deletions with MH length ≤3-nucleotide are counted as NHEJ events and >3-nucleotide are counted as aEJ events. C) deletions with MH length <4-nucleotide are counted as NHEJ events and ≤4-nucleotide are counted as aEJ events. D) deletions with MH length ≤5-nucleotide are counted as NHEJ events and >5-nucleotide are counted as aEJ events. As observed in all three illustrations, the transition point of MH lengths associated with the two processes is most likely to be at 6-to 7-nucleotide lengths. Therefore, we choose to classify NHEJ events in this study as ≤5-nucleotide MH and count MH length 6-nucleotides or longer as aEJ events for these deletions arising within the human body in normal colon crypt cells.

